# A Self-Sustaining Mechanism for Endothelial Tension Maintenance Through GqGPCR Signaling

**DOI:** 10.64898/2026.04.22.720219

**Authors:** Benjamin M. Goykadosh, Vasuretha Chandar, Harikrishnan Parameswaran

**Affiliations:** Department of Bioengineering, Northeastern University, Boston, MA, USA

**Keywords:** GqGPCR signaling, Vascular homeostasis, Mechanotransduction, Endothelial signaling

## Abstract

The vascular endothelium maintains homeostasis by acting as a selective barrier, permitting the exchange of nutrients, immune cells, and signaling molecules while restricting pathogens. It further regulates vascular function by generating and sustaining mechanical tension. Aging and disease alter the vascular environment and disrupt the regulation of endothelial tension, contributing to vascular diseases such as hypertension and atherosclerosis. Although endothelial mechanics are influenced by the cellular environment, the mechanisms that enable endothelial cells (ECs) to maintain tension over time remain poorly understood. Here, we demonstrate that confluent human umbilical vein endothelial cells (HUVECs) sustain stable tension for at least three days in the absence of external chemical or mechanical stimuli, indicating the presence of an intrinsic, active mechanism for long-term tension maintenance. Imaging of an EC multicellular ensemble shows a collective phenomenon where diacylglycerol release consistently precedes a rise in intracellular contractility. This contractility propagates to neighboring cells, wherein we identify a Gq-G-protein-coupled receptor (GqGPCR) signaling pathway as a key regulator driving force generation in ECs. The persistence of this signaling sequence in the absence of exogenous agonists suggests a ‘force-induced-force-generation’ mechanism that coordinates tension maintenance across the monolayer. Together, these findings demonstrate that ECs actively regulate tension through continuous GqGPCR signaling, revealing tension maintenance as a dynamic, collective process. This work provides new insight into how vascular tissues preserve mechanical homeostasis and suggests potential therapeutic targets for vascular endothelial dysfunction and age-related vascular stiffening.

**New and Noteworthy:** This study reveals that endothelial cells actively maintain mechanical tension through continuous GqGPCR signaling rather than passive mechanical properties. We demonstrate that DAG signaling consistently precedes contractility increases, even without chemical stimulation, suggesting that intercellular forces alone can activate this pathway. This “force-induced-force-generation” mechanism represents a potential therapeutic target for vascular dysfunction. Our findings reframe tension maintenance as a dynamic, collectively regulated process in the vascular endothelium.

## Introduction

Endothelial cells (ECs) line all blood and lymphatic vessels, maintaining tissue health and homeostasis, mediating immune cell response, and supporting angiogenesis(1–6). The endothelium acts as a barrier that selectively permits the passage of nutrients, immune cells, and signaling molecules while preventing blood leakage and protecting against pathogen invasion (2,7). A key determinant of endothelial function is its ability to generate and sustain mechanical tension which is critical for cellular processes such as barrier homeostasis, vascular remodeling and collective cell migration(8,9). Disruption of endothelial barrier integrity leads to increased permeability and dysfunction, contributing to diseases such as atherosclerosis, hypertension, fibrosis, stroke, and cancer(10–16). Endothelial tension is thought to arise from a balance between cell-cell interactions and actomyosin-based contractile forces(10). However, despite its importance, the mechanisms by which ECs generate and sustain tension under continuous hemodynamic stress remain poorly understood. Endothelial mechanics are intricately tied to their physical microenvironment. In healthy tissue, the subendothelial matrix is soft and compliant but stiffens during pathological contexts, such as aging and fibrosis(13,17,18). Increased extracellular matrix (ECM) stiffness alters endothelial behavior, whereby cells on stiffer substrates display increased cytoskeletal tension and exert larger traction forces compared to those on a softer ECM(19). Previous studies have focused on isolated ECs or cell pairs, providing limited insight into how these cells behave in their native multicellular context(20,21). In vivo, however, ECs are arranged as cohesive, mechanically integrated monolayers, in which intercellular junctions and collective force transmission underpin normal vascular function. Recent studies have further shown that ECs exhibit distinct mechanical behaviors when part of confluent monolayers compared to when isolated, highlighting the importance of multicellular context in regulating endothelial function(9,19). At the cellular level, force generation is largely attributed to actomyosin contractility regulated by intracellular calcium signaling pathways(22). Gq-G-protein–coupled receptors (GqGPCRs) are the major regulators of calcium dynamics through phosphatidylinositol-specific phospholipase C (PLC) –inositol trisphosphate (IP_3_) signaling(22). However, the role of GqGPCR signaling in coordinating and sustaining mechanical tension within confluent endothelial monolayers remains undefined. We hypothesize that intercellular mechanical forces can activate GqGPCR signaling in neighboring cells, triggering PIP2 hydrolysis and downstream calcium release in a manner analogous to agonist-induced activation. As a result, this ‘force-induced-force-generation’ mechanism would enable ECs to collectively sustain tension across a monolayer without the need for exogenous chemical stimulation. Here, we examine how ECs maintain mechanical tension within a monolayer, focusing on the interplay between substrate stiffness, cell-generated traction forces, and cell–cell interactions that regulate long-term tension maintenance. Understanding these mechanisms is fundamental to vascular biology and barrier regulation and has implications for diseases associated with endothelial mechanical dysfunction, including inflammation, atherosclerosis, and cancer(11,15,16). By elucidating how ECs sustain tension, we aim to uncover general principles of multicellular force generation and help inform studies aimed at identifying potential targets for modulating vascular function in health and disease.

## Methods and Materials

1. **Fabrication of optically clear, mechanically tunable substrates**
NuSil is an optically clear, biologically inert PDMS substrate with a tunable Young’s modulus between 0.3 to 70 kPa(23). To produce 13 kPa gels, equal parts of NuSil gel-8100, parts A and B (NuSil, CA, USA) were mixed (1:1) with a crosslinker 0.36% of total volume and spin-coated onto 30-mm-diameter, #1.5 glass coverslips for 50 seconds at 2500 RPM to produce a 100μm-thick layer. Coverslips were incubated on a flat surface at room temperature for an hour and cured overnight at 60°C. These cured coverslips were secured in sterile 40-mm Bioptech dishes (Biological Optical Technologies, Butler, PA, USA), UV-sterilized for one hour, and coated with 0.1% gelatin solution (Millipore Sigma) in preparation for use in cell culture.
2. **Endothelial Cell Culture**
Human umbilical vein endothelial cells (HUVECs) were utilized for our experiments. Cells were grown under standard cell culture conditions of 37°C and 5% CO_2_ and used before passage 15 for all experiments. Cells were cultured in Endothelial Cell Growth BBE (PSC-100-040, ATTC), 1× penicillin/streptomycin (Fisher Scientific), amphotericin B (25 μg/liter; Sigma-Aldrich). Both patterned and nonpatterned gelatin-coated substrates were UV-sterilized for 15 minutes before seeding endothelial cells. Patterns were seeded at 2.4 x 10^4 cells/mL. Cells were seeded on nonpatterned substrates at the desired density and then incubated in serum media for 24, 48 or 72 hours depending on the specific protocol. Media was then replaced with 2 mL Serum free media for 24 hours. On the day of the experiment, media was replaced with 1 mL Hanks’ Balanced Salt Solution (HBSS) (-Ca^2+^/-Mg^2+^) for 30 minutes before measurements. Nonpatterned cells were plated at the following ratios: Day 1: 525,000 cells/ dish, Day 2: 325,000 cells/ dish, Day 3: 185,000 cells/ dish for a desired confluent density of 1 x 10^6 cells/ dish at the time of measurement. All traction force (TFM) measurements were made on confluent Bioptech dishes.
3. **Traction Force Microscopy and Monolayer Stress Microscopy**
The NuSil substrates were coated with a layer of fluorescent beads to act as reference markers for TFM. A 5% solution of 0.2 µm green, fluorescent carboxylate-modified microspheres (FluoSpheres, Invitrogen, Carlsbad, CA, USA) in HBSS was vortexed for 15 seconds. A 2 mL of this mixture solution was added to each substrate in a bioptic dish and was left at room temperature on a flat surface for one hour to allow for beads to settle onto the substrate. The solution was poured off completely after one hour. Cellular tractions were recorded by imaging the fluorescent beads with a 20x/55 dry objective and the Leica DMi8 microscope. Traction force experiments were imaged at 2 second intervals for 10 minutes, with a final reference image taken after cells were removed using an RLT Lysis buffer (Qiagen, Hilden, Germany). Using these images, cellular forces were calculated using a custom MATLAB (MathWorks, Natick, MA, USA) software that utilized Fourier traction force microscopy(24).
To quantify the net contractility (M) of individual cells within a multicellular ensemble, we employed monolayer stress microscopy (MSM), a computational method that derives intracellular stresses from forces at the cell-matrix interface(25–27). The net contractility, or contractile moment, is a scalar quantity that describes the strength of the contractile dipole of a cell, and provides a single metric capturing each cell’s contractile state within the ensemble. Traction force vectors obtained from TFM were used as inputs for pyTFM, a modified open-source Python implementation of MSM, which applies a 2D in-plane force balance in accordance with Newton’s laws(28). This approach is grounded in the requirement that local traction gradients within the ensemble must be balanced by internal cellular forces. By applying a mask delineating cell-cell borders, derived from phase-contrast images of the ensemble, we resolved contractility contributions from both cell-cell and cell-matrix interactions, as well as net contractility per cell and normal and shear forces along individual cell borders. Assumptions and limitations associated with this method have been detailed previously(28,29).
4. **Matrix Protein Patterning**
In order to create cellular patterns, we used the Alvéole’s PRIMO optical module and the Leonardo software, a UV-based contactless photopatterning system (Alvéole, Paris, France). First, the NuSil surface was coated with poly-L-lysine (PLL) (500 µg/mL; Sigma-Aldrich, St. Louis, MO, USA) for 1 hour at room temperature. Then, the substrate was washed 3 times with 1X PBS and 10 mM HEPES buffer that is adjusted to pH 8.0, then incubated with methoxy polyethylene glycol-Succinimidyl Valerate (mPEG-SVA) (50 mg/ml; Laysan Bio Inc., Arab, AL, USA) at room temperature for one hour. After this point, it was washed 3 times with 1X PBS. PBS was removed and (4-benzoylbenzyl) trimethylammonium chloride (PLPP) (14.5 mg/ml; Alvéole, Paris, France) was added. The rectangular pattern (50 µm × 300 µm, H × W) was projected onto the substrate using UV illumination for 30 seconds. The surface of the gel was then washed 3 times with 1X PBS again and then incubated with 0.1% gelatin for 1 hour at room temperature. After removing the gelatin, the substrate was washed 3 times with 1X PBS and cells were plated according to desired cell density.
5. **Diacylglycerol (DAG) Transfection**
Endothelial cells were loaded with a fluorescent cytosolic diacylglycerol (DAG) indicator to record changes in DAG concentration using a live red upward diacylglycerol (DAG) assay kit (#U0350R), Montana Molecular (Montana). The assay genetically encodes fluorescent protein expression using a BacMam vector and fluorescence intensity increases with increasing DAG release upon Phosphatidylinositol 4,5-bisphosphate (PIP_2_) hydrolysis. Following the manufacturer protocol provided, we transduced cultured endothelial cells in the patterned ensemble with the DAG sensor for 24 hours at 37°C and 5% CO2. Prior to imaging, cells were washed with 1X HBSS and incubated at 37°C and 5% CO2 for an additional 30 minutes. The cells were washed once more with HBSS before imaging and transferred to serum free media prior to experiments.
6. **Data and Statistics**
Throughout the manuscript data is presented as an average of *N* independent trials. The error bars indicate the standard deviation over said independent trials. SigmaStat (Systat Software, San Jose, CA, USA) was used for calculating statistical tests. One-way ANOVAs followed by post hoc pairwise comparisons were used to test for significant differences in datasets of three or more groups, which were influenced by one independent factor. Pairwise comparisons used the *t* test when the data were normally distributed. A *P* value of 0.05 was used as the threshold to determine statistical significance between data sets.

## Results

### Endothelial cells exhibit long term tension maintenance

To elucidate how endothelial cells (ECs) sustain mechanical tension, we first characterized baseline traction forces in confluent Human umbilical vein endothelial cells (HUVEC) populations in static culture over time. Cells were seeded onto stiff (Young’s modulus E = 13 kPa (30–33)) substrates embedded with 0.2 µm fluorescent beads and cultured under standard conditions for 1, 2, and 3 days, respectively (N = 3 for each timepoint; Figure 1A). Traction forces were then quantified using traction force microscopy (TFM) over a five-minute baseline period. The confluent monolayer of ECs consistently exerted traction forces of approximately 150 Pa across all timepoints, with no statistically significant differences observed (one-way ANOVA, P = 0.98; Figure 1B). This stability in force generation suggests that HUVECs are capable of sustaining mechanical tension intrinsically, without external mechanical stretch or exogenous chemical stimulation.

**Figure 1:**
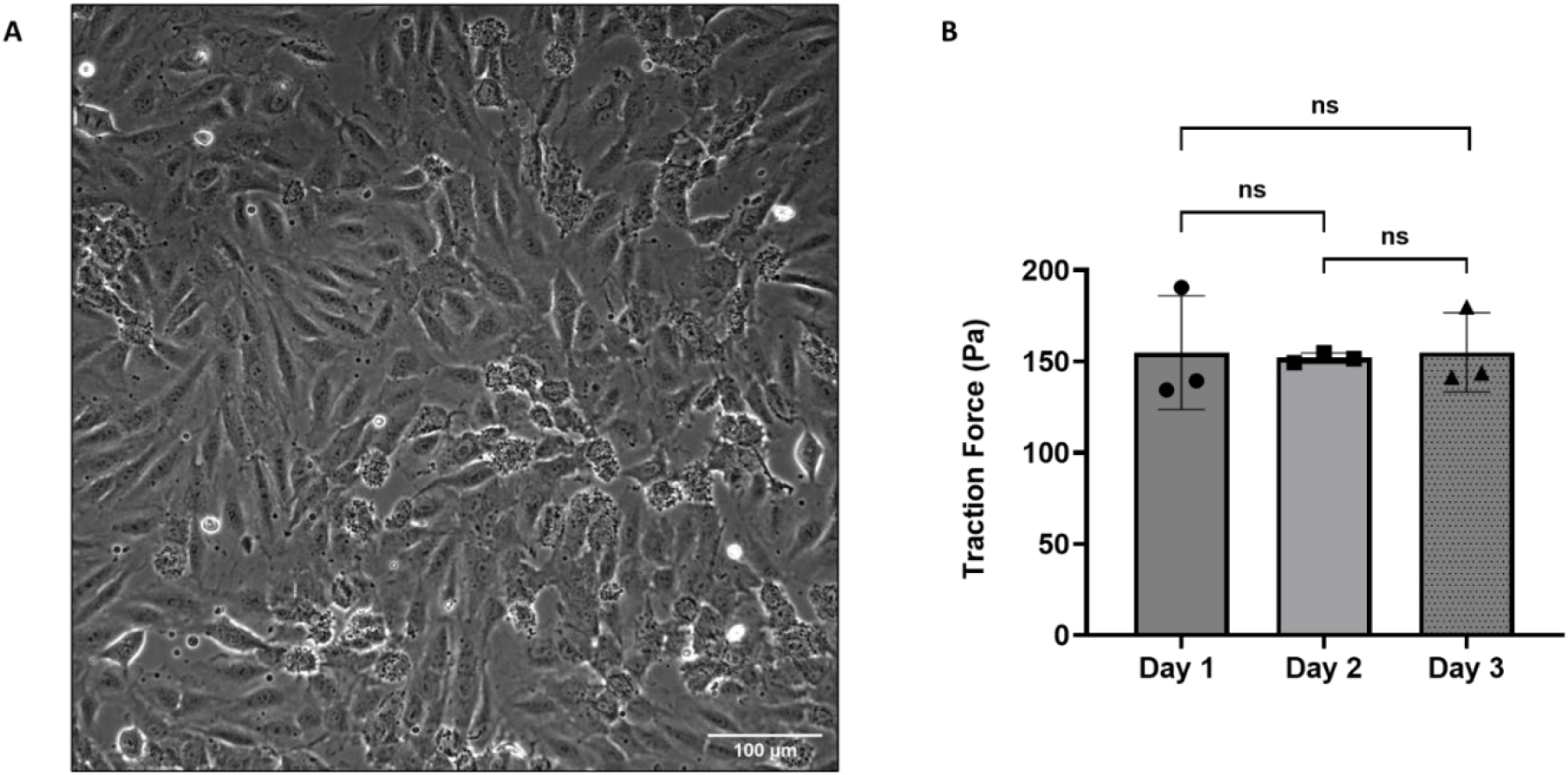
Endothelial cells can maintain tension over the course of three days. (A) <u>Representative phase image of endothelial cells</u>. Endothelial cells were seeded at confluence and cultured on 13 kPa NuSil gels embedded with fluorescent beads. (B) <u>Endothelial Cell Traction Forces Over Time</u>. HUVECs were cultured under standard conditions for one, two, and three days, and traction forces were measured at the end of each time point. Comparison of traction forces across the three timepoints demonstrated consistent force maintenance. Each data point in the bar graph refers to an individual trial (N=3). No significant difference in traction force was observed across time points (P = 0.98, one-way ANOVA), indicating that endothelial cells are able to intrinsically maintain traction forces and tension over three days.

### Endothelial cells maintain tension via the GqGPCR pathway

To investigate how ECs maintain mechanical tension in confluent ensembles, we examined a molecular pathway previously identified by Berridge et al. (22). There, it was established that activation of a Gq-G-protein-coupled receptor (GqGPCR) by an agonist initiates a signaling cascade that cleaves phosphatidylinositol 4,5-bisphosphate (PIP_2_) into diacylglycerol (DAG) and inositol 1,4,5-trisphosphate (IP_3_). IP_3_ triggers intracellular Ca^2+^ release, which in turn drives force generation and maintenance.

Building on this, we investigated the temporal relationship between DAG signaling and contractility (M) in ECs to explore whether a similar regulatory mechanism exists where intercellular forces can activate the GqGPCR pathway (Figure 2.A).

**Figure 2:**
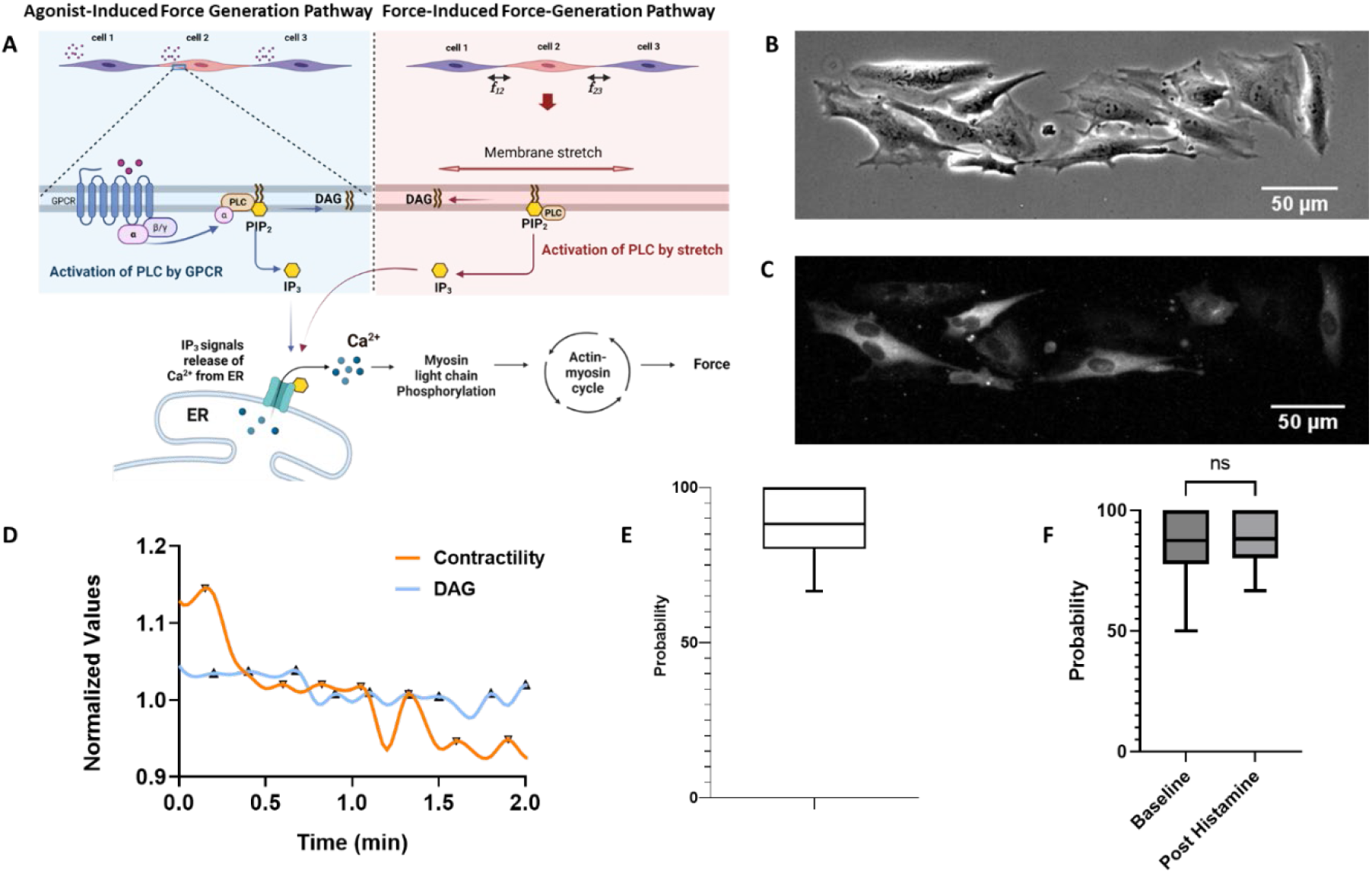
(A) Schematic of a force-based force generation mechanism. We hypothesize that HUVECs can use their force to modulate force maintenance in neighboring HUVECs. Just as an agonist binding to a GqGPCR results in PIP_2_ hydrolysis to form IP_3_ **(Right panel)**, intercellular force communication can hydrolyze PIP_2_ to form DAG and IP_3_ **(Left Panel)**. This IP_3_ release in turn binds to receptors on the ER and release Ca^2+^, setting off the same chain of reactions that lead to force generation. (B) Representative phase image of a patterned endothelial ensemble of cells cultured on a 13kPa stiff substrate. Scale bar 50 um. (C) Representative fluorescence image showing intracellular DAG elevation after agonist addition, where white fluorescence indicates DAG signaling. 12 positively transfected cells out of 14 cells in the ensemble exhibited a rapid increase in DAG signaling, indicated by increased fluorescence intensity, corresponding to activation and initiation of cellular contraction. (D) Representative time series of DAG (blue) and contractility (M, orange) peaks for a single cell, demonstrating that a peak in DAG consistently precedes a peak in contractility following GqGPCR pathway activation. (E) Distribution of conditional probability of the hypothesized signaling sequence (DAG peak between M peaks) post-histamine treatment. Out of a total of 21 positively transfected cells across 3 trails, the conditional probability was 100% for 6 cells, 80–100% for 10 cells, and 65– 80% for 5 cells. (F) Comparison of conditional probability of the hypothesized signaling sequence (DAG peak between M peaks) at baseline and post-histamine treatment. No significant difference was observed (P = 0.83, paired t-test, N = 3 trials, 21 cells total), indicating that mechanical forces contribute to tension maintenance independent of chemical stimulation.

To this end, HUVECs were transfected to express fluorescently tagged DAG markers and were geometrically confined into rectangular ensembles of positively transfected cells on 13 kPa NuSil gels embedded with fluorescent beads. Geometric confinement defined the monolayer boundary, ensuring reproducible cell-cell contact and enabling accurate computation of intercellular stresses. Cells were imaged 24 hours post seeding. M and DAG values were recorded every two seconds for five minutes to establish baseline activity, after which cells were stimulated with 10^−5^ M histamine (Figure 2.B-C). Only successfully transfected cells were included in the analysis (N = 3 independent trials, 21 cells positively transfected).

Net contractility of individual HUVECs was quantified using an open-source implementation of Monolayer Stress Microscopy (MSM)(28,29). This tool computes in-plane mechanical stresses within the monolayer, producing detailed maps of intercellular tension and enabling independent measurement of cell-cell and cell-matrix contractile forces(24–27). The contractile moment, or contractility, is a scalar quantity which indicates the strength of the contractile dipole of a cell.

Using this data, we analyzed the conditional probability that a local maximum in DAG signaling occurred between two local maxima in the M signal. If DAG and M dynamics were independent, DAG peaks would be randomly distributed relative to M peaks, yielding a conditional probability near 50%. Across three independent experiments, DAG peaks consistently preceded peaks in contractility (Figure 2.D). Out of the 21 positively transfected cells across three trials, the proposed M-DAG sequence occurred 100% of the time in 6 cells, 80–100% in 10 cells, and 65–80% in 5 cells. These findings support a model in which GqGPCR signaling and DAG production are critical for the generation and maintenance of contractile force in ECs (Figure 2.E). Importantly, this signaling sequence was preserved both before and after histamine stimulation (Figure 2.F), suggesting that the M-DAG coordination is not only involved in the acute response to stimulation but also plays a sustained role in basal tension maintenance. Together, these findings indicate that ECs possess an intrinsic mechanism for maintaining mechanical tension through sustained cytoskeletal engagement.

## Discussion

In this study, we investigated how endothelial cells (ECs) maintain mechanical tension as confluent ensembles. We found that HUVECs sustain consistent traction forces of approximately 150 Pa over three days in culture, with no significant changes between timepoints. This stability demonstrates that ECs possess an intrinsic capacity for long-term tension maintenance. To explore the molecular basis of this behavior, we examined the temporal relationship between DAG signaling and contractility (M). We found that DAG peaks consistently preceded M peaks, and importantly, this signaling sequence persisted both before and after histamine stimulation. These findings suggest that GqGPCR pathway activation underlies active force maintenance in ECs and that this pathway can be engaged by mechanical cues independent of exogenous agonists. This process is actively regulated through sustained signaling rather than being a passive consequence of cytoskeletal properties.

Our observation that the M-DAG signaling sequence occurs under baseline conditions without chemical stimulation suggests that intercellular mechanical forces can activate the GqGPCR pathway. To quantify these forces, we employed Monolayer Stress Microscopy, which enabled us to compute intercellular stresses within the monolayer and distinguish between cell-cell and cell-matrix contractile contributions.

This approach was essential for testing our hypothesis that force communication between neighboring cells drives pathway activation. The consistency of the M-DAG sequence across cells points to a self-sustaining feedback mechanism in which force generation by one cell mechanically stimulates signaling in neighboring cells, thereby coordinating tension maintenance across the ensemble. Consistent with this model, emerging evidence suggests that PLC can be activated by mechanical stimuli. Mechanical activation of GqGPCRs has been shown to drive PLC-mediated PIP_2_ hydrolysis, and membrane stretch alone is sufficient to induce PLC-dependent PIP_2_ hydrolysis(22,34–36). These findings raise the possibility that intercellular forces transmitted through cell-cell junctions may similarly engage PLC activity, providing a mechanistic link between force transmission and the DAG signaling we observe.

Such a mechanism would allow endothelial monolayers to maintain barrier integrity and respond dynamically to changes in hemodynamic load.

It is important to note that our work was performed under static conditions to isolate the intrinsic force maintenance capabilities independent of external mechanical stimulation. In vivo, however, the endothelium is continuously exposed to fluid shear stress and under those conditions ECs align in the direction of flow(37). This uniform orientation is likely to influence the distribution of intercellular stresses and, by extension, the M-DAG signaling dynamics we describe. Additionally, experiments were conducted on a single substrate stiffness, and it remains to be determined how varying matrix compliance influences these dynamics. However, these limitations point toward productive future directions incorporating pharmacological inhibitors or genetic knockdown approaches. This would further help define the mechanistic link between GqGPCR signaling and force maintenance, and to identify specific molecular targets for therapeutic intervention. By establishing that endothelial tension maintenance is an actively regulated signaling process, this work provides a framework for understanding vascular mechanical homeostasis. Understanding how the endothelium maintains tension may ultimately inform therapeutic strategies for conditions characterized by endothelial dysfunction, including hypertension, fibrosis, and cancer.

## Author Contributions

H.P. conceived the research and developed the overall hypothesis, and B.M.G and H.P. developed the experimental design. B.M.G. conducted the experiments, analyzed the data, and interpreted the results along with H.P. B.M.G. prepared the figures. B.M.G and V.C. conducted the Monolayer Stress Microscopy experiments. B.M.G. analyzed the monolayer stress microscopy data, prepared the figures and interpreted the results. B.M.G, V.C. and H.P contributed to the drafting and editing of the manuscript. H.P approved the final version of the manuscript.

